# SpatialMOC: Accurate reconstruction of spatial multi-omics landscapes through cross-modality prediction

**DOI:** 10.64898/2026.08.10.743892

**Authors:** Zhengxuan Liu, Peimeng Zhen, Bingtao Wang, Han Shu, Yuan Zhao, Kaiyue Zhang, Yongtian Wang, Jialu Hu, Tao Wang, Jing Chen

## Abstract

Spatial multi-omics technologies provide unprecedented opportunities to characterize tissue organization by measuring complementary molecular layers within their native spatial context. However, simultaneous profiling of multiple molecular modalities remains technically challenging, limiting the widespread application of spatial multi-omics and leaving most studies reliant on single-modality measurements. Here, we present Spatial Multi-Omics Cross-prediction (Spatial-MOC), a computational framework that reconstructs missing spatial molecular modalities by integrating spatial context with cross-modality representation learning. Across multiple tissues, molecular modalities, developmental stages, sequencing platforms, and degraded datasets, SpatialMOC consistently outper-formed existing computational approaches in bidirectional molecular prediction. Beyond accurate prediction, SpatialMOC faithfully reconstructed tissue architecture, preserved dynamic molecular and regulatory heterogeneity, and recovered biologically meaningful spatial landscapes from technically compromised measurements. Together, these results establish SpatialMOC as a general framework for spatial multi-omics reconstruction, extending the analytical value of existing spatial omics datasets and facilitating comprehensive investigations of tissue organization, development, and disease.

## Introduction

Spatially resolved transcriptomics and related multi-omics technologies have transformed the study of tissue organization by mapping molecular measurements to their native anatomical context [1, 2]. Joint measurements of gene expression with either chromatin accessibility or protein abundance can connect cellular states to regulatory mechanisms and tissue microenvironments [3–5]. Nevertheless, collecting several molecular layers from the same tissue remains constrained by technical complexity, targeted or incomplete molecular coverage, and modality-specific noise or sparsity [6, 7]. These advances motivate extending missing-modality prediction from dissociated single cells to spatially resolved multi-omics data [8, 9].

Several computational frameworks have been developed for cross-modality prediction and multimodal integration in dissociated single-cell data. Encoder-decoder architectures, including BABEL [10], CMAE [11], scMOG [12], and scPair [13], directly learn nonlinear mappings between molecular modalities. Probabilistic generative models such as MultiVI [14] and totalVI [15] jointly model paired transcriptomic, chromatin accessibility, or protein measurements, whereas graph-based approaches, including scMoGNN [16], explicitly capture relationships among cells. Complementary integration methods, including Seurat [17, 18], MOFA+ [19], and LIGER [20], align multiple molecular modalities through shared latent representations or linked feature spaces. Despite their success, most methods do not explicitly incorporate spatial coordinates to preserve tissue architecture during cross-modality prediction.

Recent advances have extended computational modeling to spatial multi-omics data. SpatialGlue [21] and dependency-aware spatial generative models exploit spatial neighborhoods and cross-omics dependencies to improve representation learning and spatial domain identification. LLOKI [22] addresses cross-platform integration of imaging-based spatial transcriptomics, whereas EYKTHYR [23] integrates spatial transcriptomic and chromatin accessibility data to infer spatial gene regulatory programs. More recently, SpaMIE [24] introduced a graph-based framework for missing-modality imputation under heterogeneous modality coverage. However, existing methods primarily focus on modality completion or integration and generally do not provide a unified framework for bidirectional cross-modality prediction while simultaneously preserving tissue architecture across diverse spatial multi-omics datasets.

To address these challenges, we developed Spatial Multi-Omics Cross-prediction (SpatialMOC), a computational framework for bidirectional prediction and reconstruction of spatial multi-omics profiles. SpatialMOC jointly models spatial context and cross-modality correspondence by combining spatially constrained latent representations with probabilistic generative modeling, enabling accurate molecular prediction while preserving tissue architecture. A shared latent space aligns heterogeneous molecular modalities for robust cross-omics translation, and modality-specific observation models accommodate the distinct statistical properties of transcriptomic, chromatin accessibility, and proteomic measurements. Together, these components provide a unified framework for spatial multi-omics prediction across diverse molecular modalities and experimental settings.

We evaluated SpatialMOC using paired RNA, ATAC, and ADT measurements generated by MISAR-seq, SPOTS, Spatial-CITE-seq, and spatial epigenome-transcriptome co-profiling across multiple tissues, developmental stages, sequencing platforms, and data quality conditions. Benchmarking included bidirectional RNA–ATAC and RNA–ADT prediction, assessment of spatial domain preservation, and evaluation under cross-stage, cross-platform, and simulated-dropout settings. These experiments demonstrate the robustness and generalizability of SpatialMOC for spatial multi-omics prediction across diverse biological and technical scenarios.

## Methods

### Spatial Multi-omics Input and Preprocessing

#### Cross-omics dependency modeling

Consider a paired spatial multi-omics dataset comprising *N* spatial locations with two-dimensional coordinates *X* ∈ ℝ^*N* ×2^. Two molecular modalities are measured at each location and represented by count matrices 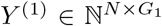 and 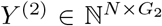, where *G*_1_ and *G*_2_ denote the numbers of modality-specific features.

SpatialMOC assumes that both molecular modalities are generated from a shared latent representation *Z* ∈ ℝ^*N* ×*D*^, where *D* ≪ min(*G*_1_, *G*_2_), following the general latent-variable formulation used in deep generative models of molecular count data [25]. The joint likelihood of the observed data is defined as

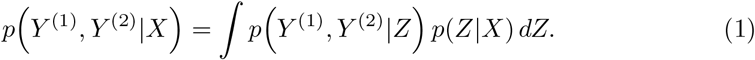

Assuming conditional independence between the two molecular modalities given the latent representation, the likelihood factorizes as

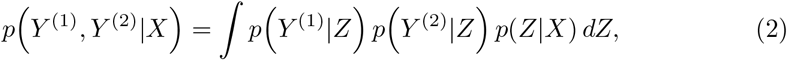

where *p*(*Z*|*X*) denotes a spatial prior conditioned on tissue coordinates. Because direct posterior inference is intractable, SpatialMOC employs variational inference to approximate the latent distribution [26, 27], as described in the following sections.

#### Unified data preprocessing pipeline

All spatial multi-omics datasets were preprocessed using Scanpy [28] and Squidpy. For spatial transcriptomics (RNA), low-quality spots containing fewer than 100 unique molecular identifiers (UMIs), fewer than 50 detected genes, or more than 20% mito-chondrial transcripts were removed. The remaining data were library-size normalized and log-transformed, after which the top 3,000 highly variable or spatially variable genes were selected for downstream analyses.

For ATAC data, low-quality spots and low-frequency peaks were removed before term frequency–inverse document frequency (TF–IDF) normalization. The top 3,000– 5,000 highly variable peaks were retained according to TF–IDF scores and converted into a binary accessibility matrix for model training.

For spatial proteomics, ADT counts were normalized using the centered log-ratio (CLR) transformation.

Finally, two-dimensional spatial coordinates were independently standardized along each axis to zero mean and unit variance before spatial graph construction and model training.

### Spatial Variational Encoder

To learn a shared latent representation while preserving tissue architecture, Spatial-MOC employs an omics-specific spatial variational encoder. Each encoder decomposes the latent representation into spatial and non-spatial components and incorporates a Gaussian process prior to model spatial dependencies among neighboring locations.

#### Latent space factorization

For each molecular modality *m* ∈ {1, 2}, the latent representation *Z*^(*m*)^ ∈ ℝ^*N* ×*D*^ is partitioned into a spatial component 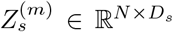 and a non-spatial component 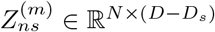,

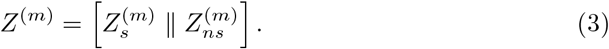

Assuming conditional independence between the two latent components given the observed molecular measurements, the variational posterior is factorized as

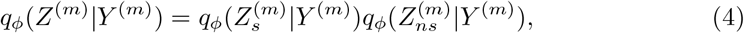

where *ϕ* denotes the parameters of the omics-specific encoder.

The non-spatial latent variables are modeled using a multivariate Gaussian distribution,

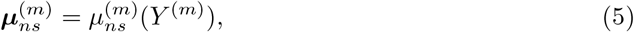

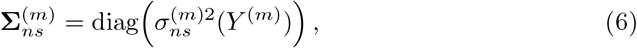

yielding the approximate posterior

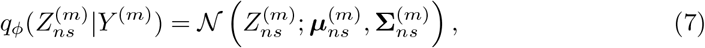

which is regularized by an isotropic Gaussian prior,

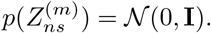

#### Shared spatial Gaussian process prior

The spatial latent variables are constrained by a Gaussian process (GP) prior,

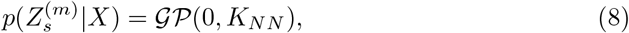

where *K*_*NN*_ denotes the covariance matrix evaluated at the observed spatial coordinates *X*.

To ensure that the latent representations inferred from different molecular modalities preserve a common tissue architecture, all modality-specific encoders share the same spatial covariance function. The covariance between two spatial locations *x*_*i*_ and *x*_*j*_ is computed using the Cauchy kernel,

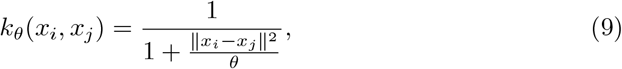

where *θ* is a learnable scale parameter. Compared with the squared-exponential kernel, the heavier-tailed Cauchy kernel retains correlations over longer spatial distances.

The spatial latent variables are regularized by minimizing the Kullback–Leibler (KL) divergence between the variational posterior and the GP prior.

#### Sparse variational Gaussian process approximation

Exact Gaussian process inference requires inversion of the dense covariance matrix *K*_*NN*_, resulting in a computational complexity of *O*(*N* ^3^) and memory complexity of *O*(*N* ^2^).

To enable scalable training, SpatialMOC adopts a sparse variational Gaussian process (SVGP) approximation [29]. A set of *P* inducing points located at pseudo-coordinates *X*_*p*_ (*P* ≪ *N*) is introduced to approximate the full GP prior. Conditioning the latent representation on the inducing variables reduces the computational complexity to *O*(*bP* ^2^) for a mini-batch of size *b*, enabling efficient optimization on high-resolution spatial multi-omics datasets while preserving the underlying spatial dependency structure.

### Cross-modality Latent Alignment

To obtain a shared latent representation across molecular modalities, SpatialMOC employs an adversarial alignment strategy. A discriminator network *D*_*ψ*_ is trained to identify the modality of origin for each latent representation, whereas the omics-specific encoders 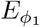 and 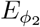 are optimized to generate modality-invariant embeddings. Consequently, the latent distributions satisfy

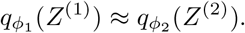

The adversarial objective is defined as

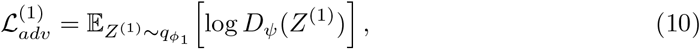

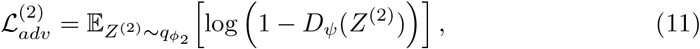

yielding the minimax optimization problem

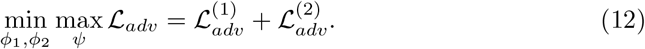

Under the optimal discriminator, this objective corresponds to minimizing a divergence between the modality-specific latent distributions that is closely related to the Jensen–Shannon divergence [30].

During optimization, the discriminator maximizes the adversarial objective to distinguish the two modalities, whereas the encoders minimize the objective to learn a common latent representation suitable for cross-modality prediction.

### Omics-specific Generative Decoder

The aligned latent representations are decoded into modality-specific observation spaces using likelihood models that match the statistical characteristics of each molecular modality.

#### Transcriptomics (RNA)

Spatial transcriptomic counts are modeled using a negative binomial (NB) distribution [31],

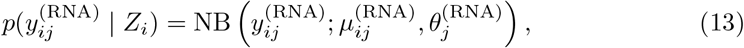

where the mean parameter is decomposed as

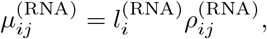

with 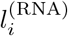 denoting the empirical library size and 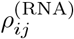 representing the normalized gene expression predicted by the decoder.

#### Chromatin accessibility (ATAC)

Spatial chromatin accessibility is modeled using a Bernoulli distribution,

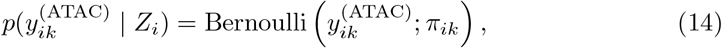

where the accessibility probability is given by

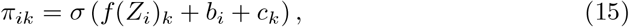

where *f* (·) denotes the decoder network, *b*_*i*_ is a spot-specific bias, and *c*_*k*_ is a peak-specific bias.

#### Spatial proteomics (ADT)

Spatial protein counts are modeled using a negative binomial distribution,

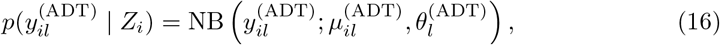

where 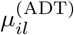 and 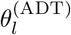 denote the decoder-predicted mean and protein-specific dispersion parameter, respectively.

### Model Optimization

Model parameters are optimized by maximizing the evidence lower bound (ELBO), which jointly optimizes the modality-specific reconstruction likelihood and the variational regularization imposed on the latent representations.

For each molecular modality *m*, the ELBO is defined as

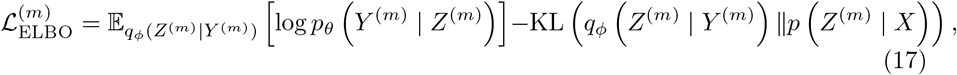

where the first term corresponds to the modality-specific reconstruction likelihood and the second term regularizes the latent representation using the spatial Gaussian process prior.

The reconstruction loss is defined as the negative sum of the modality-specific ELBOs,

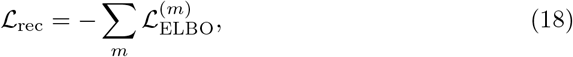

where the negative ELBO is minimized during optimization.

The final objective combines the reconstruction loss with the adversarial alignment loss,

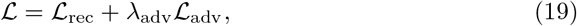

where *λ*_adv_ controls the contribution of adversarial latent alignment.

Model parameters were optimized using the Adam optimizer with mini-batch stochastic gradient descent [32]. During each iteration, the discriminator and encoder networks were updated alternately until convergence.

### Clustering

Spatial domains were identified by applying *k*-means clustering to the integrated latent representations learned by SpatialMOC. For all datasets, the number of clusters (*k*) was set according to the annotated anatomical structures to facilitate quantitative evaluation. Specifically, *k* was set to 9 for the mouse brain, 8 for the human tonsil, 6 for the mouse thymus, and 5 for the mouse spleen.

### Benchmark

SpatialMOC was compared with seven baseline methods: SpaMIE, BABEL, the CMAE framework, scPair, MultiVI, totalVI, and scMoGNN. For each modality pair, the applicable labels and values are reported in the Supplementary Tables.

Prediction performance was evaluated using the Pearson correlation coefficient (PCC) for continuous RNA and ADT prediction and for ATAC-derived gene-activity scores. The area under the receiver operating characteristic curve (AUROC) was used when evaluating binary chromatin accessibility. Sample-level correspondence between predicted and observed target-modality profiles was evaluated using one minus the fraction of samples closer than the true match (1 − FOSCTTM), where a larger value indicates better correspondence[33]. Spatial domain identification was assessed using the adjusted Rand index (ARI) [34], normalized mutual information (NMI), adjusted mutual information (AMI) [35], and homogeneity score (HOMO) [36]. Additional implementation details are provided in the Supplementary Information.

### Data availability

The spatial cellular indexing of transcriptomes and epitopes by sequencing (Spatial-CITE-seq) human tonsil and mouse spleen datasets were obtained from the Gene Expression Omnibus (GEO) under accession GSE213264. The Spatial PrOtein and Transcriptome Sequencing (SPOTS) mouse spleen dataset is available under GSE198353. The spatial ATAC–RNA mouse brain dataset was obtained from AtlasX-plore. The unpublished Spatial Enhanced Resolution Omics Sequencing (Stereo-seq) combined with CITE-seq (Stereo-CITE-seq) mouse thymus datasets are available in the SpatialGlue dataset record on Zenodo (version v3) [37]. The Microfluidic Indexing-based Spatial Assay for chromatin accessibility and RNA sequencing (MISAR-seq) mouse embryo datasets were obtained from the National Genomics Data Center under accession OEP003285.

## Results

### SpatialMOC enhances spatial multi-omics prediction through spatially aware representation learning

SpatialMOC is a computational framework designed to predict missing molecular modalities from spatial multi-omics data while preserving the spatial organization of biological tissues (Figure 1A). The framework addresses two major challenges in spatial multi-omics prediction: accurately modeling spatial dependencies within tissues and enabling generalizable cross-modality prediction across heterogeneous molecular profiles. To achieve this, SpatialMOC takes paired spatial multi-omics data from a reference tissue together with incomplete spatial measurements from a target tissue as input and reconstructs the missing molecular modalities within a unified prediction framework.

**Fig. 1.**
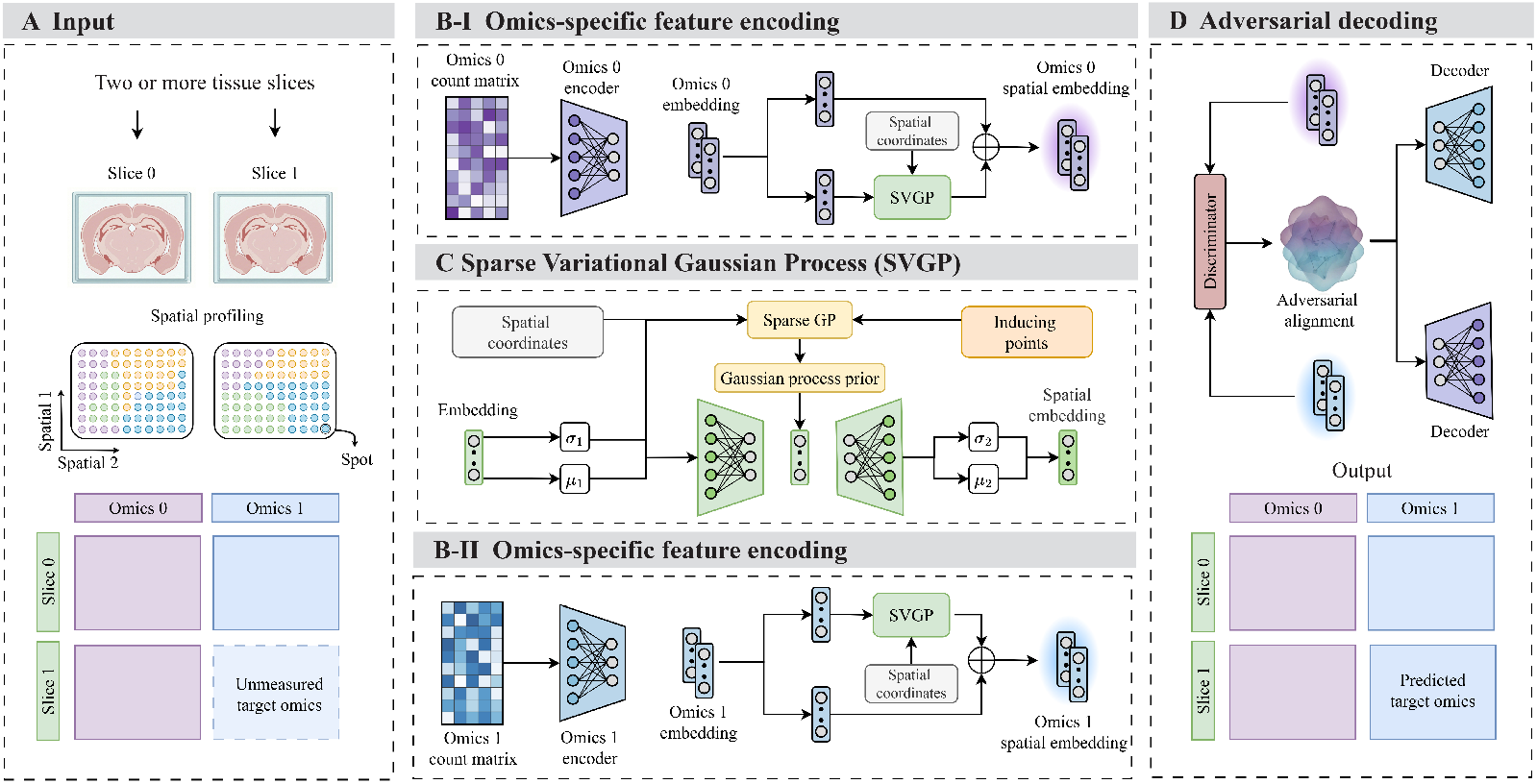
Overview of the SpatialMOC framework. **A**, Overall workflow of SpatialMOC for spatial multi-omics prediction. The framework takes paired multi-omics data from a reference tissue together with incomplete measurements from a target tissue and reconstructs the missing molecular modalities while preserving spatial organization. **B**, Omics-specific representation learning. Dual-branch encoders learn complementary molecular and spatial representations from each omics, enabling effective integration of molecular features with spatial context. **C**, Spatial dependency modeling. A sparse variational Gaussian process (SVGP) introduces a shared spatial prior that captures continuous spatial relationships and preserves tissue architecture across molecular modalities. **D**, Cross-modality alignment and prediction. An adversarial representation learning module aligns heterogeneous omics into a shared latent space, after which omics-specific probabilistic decoders reconstruct the target molecular modalities for accurate cross-omics prediction and spatial interpolation.

SpatialMOC consists of three complementary components that jointly support robust cross-modality prediction (Figure 1B–D). First, an omics-specific representation learning module extracts latent molecular features while explicitly incorporating spatial information, thereby preserving local tissue architecture and spatial molecular patterns. Second, a spatially aware variational learning strategy based on a sparse variational Gaussian process (SVGP) models continuous spatial dependencies by introducing a shared spatial prior across molecular modalities. Finally, an adversarial representation alignment module projects heterogeneous omics into a common latent space, enabling efficient knowledge transfer between modalities while reducing modality-specific biases. The aligned latent representations are subsequently decoded using omics-specific probabilistic models to reconstruct the target molecular profiles.

By integrating spatial representation learning, cross-modality alignment, and distribution-aware decoding within a unified framework, SpatialMOC provides accurate and flexible prediction across diverse spatial multi-omics datasets. The framework is applicable to multiple molecular modalities and experimental settings, supports downstream analyses such as spatial domain identification and spatial molecular interpolation, and remains robust to heterogeneous data quality. These architectural features establish the foundation for the comprehensive benchmarking and biological evaluations presented in the following sections.

### SpatialMOC enables accurate bidirectional prediction across multiple spatial molecular modalities

To evaluate the ability of SpatialMOC to reconstruct missing molecular modalities from spatial single-omics measurements, we performed cross-slice prediction experiments using two complementary spatial multi-omics datasets: the MISAR-seq mouse embryo dataset at embryonic day 15.5 (E15.5), comprising paired spatial RNA and ATAC measurements, and the SPOTS mouse spleen dataset, comprising paired spatial RNA and ADT profiles. In each experiment, SpatialMOC was trained on a reference tissue section with paired multi-omics data and subsequently used to predict the missing molecular modality in an adjacent target section.

We first assessed bidirectional prediction between spatial RNA and ATAC using adjacent sections from the mouse embryo E15.5 dataset, in which ATAC profiles were converted to gene activity scores for quantitative evaluation. The anatomical annotations of the reference tissue are shown in Figure 2A. SpatialMOC consistently achieved the highest prediction accuracy among all methods, reaching Pearson correlation coefficient (PCC) values of 0.8412 for RNA-to-ATAC prediction and 0.8221 for ATAC-to-RNA prediction, followed by SpaMIE (0.7722 and 0.7288) and scPair (0.7443 and 0.7012) (Supplementary Tables S1 and S2). The overall comparison is shown in Figure 2B. Similar improvements were observed for the 1 − FOSCTTM metric, with median scores of 0.9133 and 0.8871 for the two prediction tasks, respectively (Figure 2C). Complete quantitative comparisons are provided in Supplementary Tables S1 and S2.

**Fig. 2.**
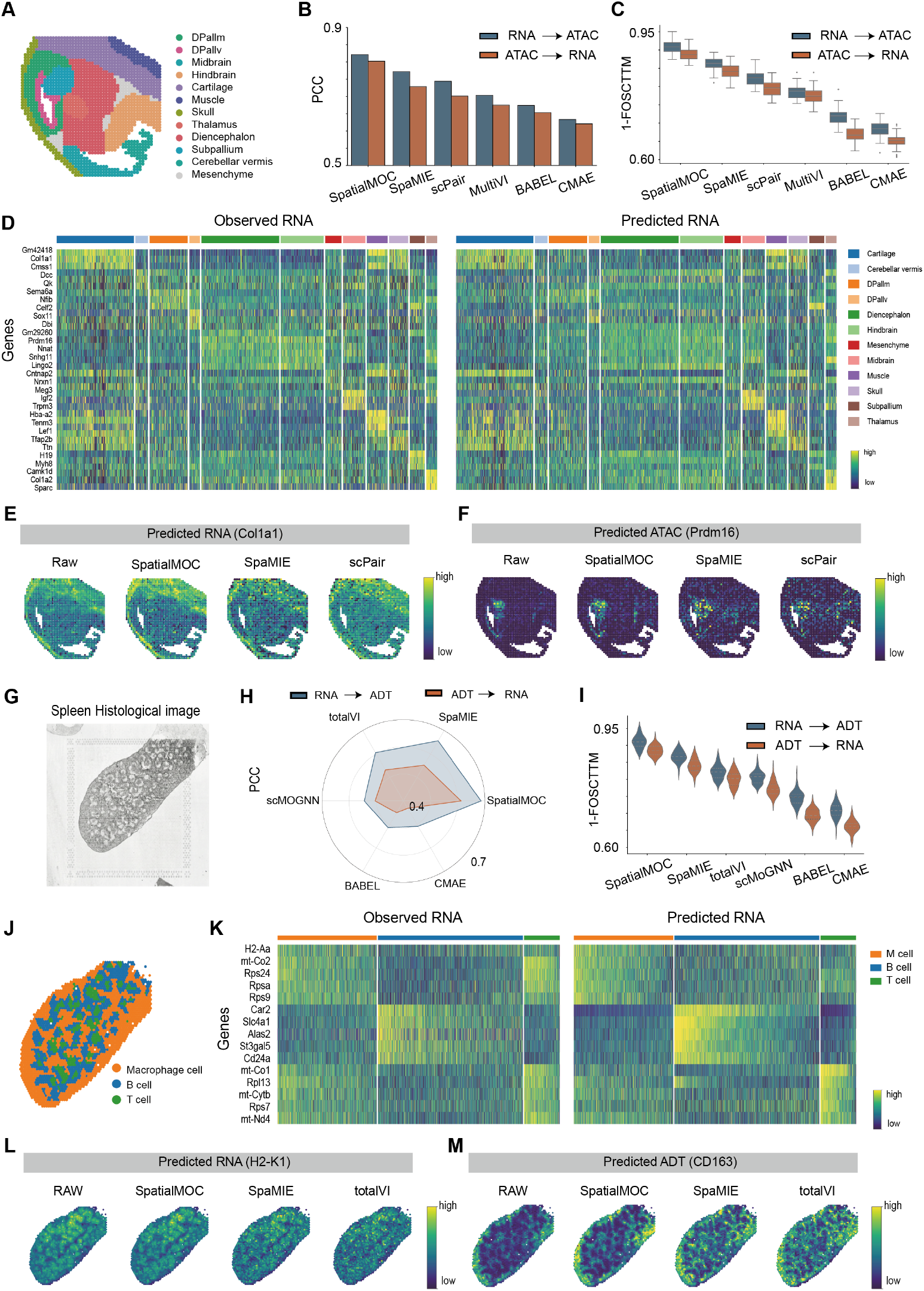
SpatialMOC accurately reconstructs diverse spatial molecular modalities across tissues. (**A**) Anatomical annotations of the embryonic day 15.5 (E15.5) mouse embryo used for RNA–ATAC prediction. (**B**) Quantitative comparison of bidirectional RNA-to-ATAC and ATAC-to-RNA prediction accuracy measured by PCC. (**C**) Comparison of prediction performance using the 1 − FOSCTTM metric for bidirectional RNA–ATAC prediction. (**D**) Heatmap comparing observed and predicted RNA expression across annotated anatomical regions. (**E**) Spatial reconstruction of the representative RNA marker *Col1a1*, compared with the observed data and baseline methods. (**F**) Spatial reconstruction of the representative ATAC marker *Prdm16*. (**G**) Histological image of the mouse spleen used for RNA–ADT prediction. (**H**) Quantitative comparison of bidirectional RNA-to-ADT and ADT-to-RNA prediction accuracy measured by PCC. (**I**) Comparison of prediction performance using the 1 − FOSCTTM metric for bidirectional RNA–ADT prediction. (**J**) Spatial distribution of immune cell clusters in the mouse spleen. (**K**) Heatmap comparing observed and predicted RNA expression across immune cell clusters. (**L**) Spatial reconstruction of the representative RNA marker *H2-K1*, compared with the observed data and baseline methods. (**M**) Spatial reconstruction of the representative ADT marker CD163.

To further assess the biological fidelity of the predicted profiles, we compared reconstructed molecular landscapes with the observed spatial data. The reconstructed transcriptomic heatmaps closely recapitulated the observed expression patterns across anatomical regions (Figure 2D). Consistently, SpatialMOC accurately recovered the spatial distribution of representative molecular markers (Figure 2E–F). In particular, the predicted RNA expression of *Col1a1* faithfully reproduced its enrichment within cartilage, muscle, and skull regions, whereas the predicted ATAC activity of *Prdm16* correctly localized to the subpallium and closely matched the observed spatial accessibility pattern. These results demonstrate that SpatialMOC accurately reconstructs both transcriptomic and chromatin accessibility landscapes while preserving biologically meaningful spatial organization.

We next evaluated the generalizability of SpatialMOC using the SPOTS mouse spleen dataset, which contains paired spatial RNA and ADT measurements (Figure 2G). Despite the distinct molecular modalities and tissue architecture, SpatialMOC consistently maintained superior prediction performance. It achieved the highest PCC values of 0.6859 for RNA-to-ADT prediction and 0.6123 for ADT-to-RNA prediction while simultaneously obtaining the best 1 − FOSCTTM scores (0.8147 and 0.7845), followed by SpaMIE, totalVI, and other competing methods (Supplementary Tables S3 and S4). The overall comparison is shown in Figure 2H,I. Detailed quantitative results are summarized in Supplementary Tables S3 and S4.

Beyond quantitative performance, SpatialMOC faithfully preserved the spatial organization of immune cell populations in the spleen. Immune-cell annotations are shown in Figure 2J. The reconstructed transcriptomic heatmaps accurately recapitulated the observed molecular patterns across immune cell clusters identified by *k*-means clustering (Figure 2K). Furthermore, the predicted spatial expression of representative markers closely matched the experimental observations (Figure 2L–M). The predicted expression of the major histocompatibility complex (MHC) class I gene *H2-K1* reproduced its characteristic nodular enrichment within the white pulp, whereas the predicted distribution of the macrophage marker CD163 accurately delineated the reticular architecture of the red pulp. Notably, these two complementary spatial patterns were faithfully reconstructed despite being inferred from different molecular modalities, indicating that SpatialMOC preserves higher-order tissue compartmentalization and biologically meaningful tissue architecture during cross-modality prediction.

Finally, to further evaluate the robustness of SpatialMOC, we performed reciprocal prediction analyses by reversing the reference and target tissue sections in both the MISAR-seq and SPOTS datasets. SpatialMOC consistently achieved the best performance across all evaluation metrics regardless of prediction direction. Detailed quantitative results and corresponding visualizations are provided in Supplementary Tables S5–S8 and Supplementary Figures S1–S4.

### SpatialMOC faithfully reconstructs tissue architecture from predicted spatial molecular profiles

To determine whether SpatialMOC preserves biologically meaningful tissue architecture during cross-modality prediction, we evaluated its ability to reconstruct spatial domains using two complementary datasets: a spatial ATAC–RNA mouse brain dataset and a Spatial-CITE-seq human tonsil dataset. Because accurate spatial domain identification depends on the preservation of local molecular organization, these analyses provide a stringent assessment of structural fidelity following cross-omics prediction. We performed five-fold cross-validation, in which one subset of spatial spots was withheld for prediction in each iteration. Predicted molecular profiles from all folds were subsequently combined to reconstruct complete tissue-wide molecular maps for downstream spatial clustering and evaluation.

We first evaluated bidirectional RNA–ATAC prediction using the mouse brain dataset. The anatomical annotations of the reference tissue are shown in Figure 3A, highlighting distinct cortical layers (L1–3, L4, L5, and L6a/b) together with subcortical structures including the caudoputamen (CP) and nucleus accumbens (ACB). Quantitative evaluation demonstrated that SpatialMOC consistently achieved the highest structural consistency across all clustering metrics. In the RNA-to-ATAC prediction task, SpatialMOC obtained an ARI of 0.8128 and an NMI of 0.8328, followed by SpaMIE (ARI: 0.7323; NMI: 0.7495) (Supplementary Table S9). Similar improvements were observed for ATAC-to-RNA prediction, where SpatialMOC achieved an ARI of 0.8028 compared with 0.6815 for SpaMIE (Supplementary Table S10). The overall comparisons are shown in Figure 3B,C. Complete quantitative results for all five cross-validation folds are provided in Supplementary Tables S9 and S10, with corresponding distributions shown in Supplementary Figures S5–S8.

**Fig. 3.**
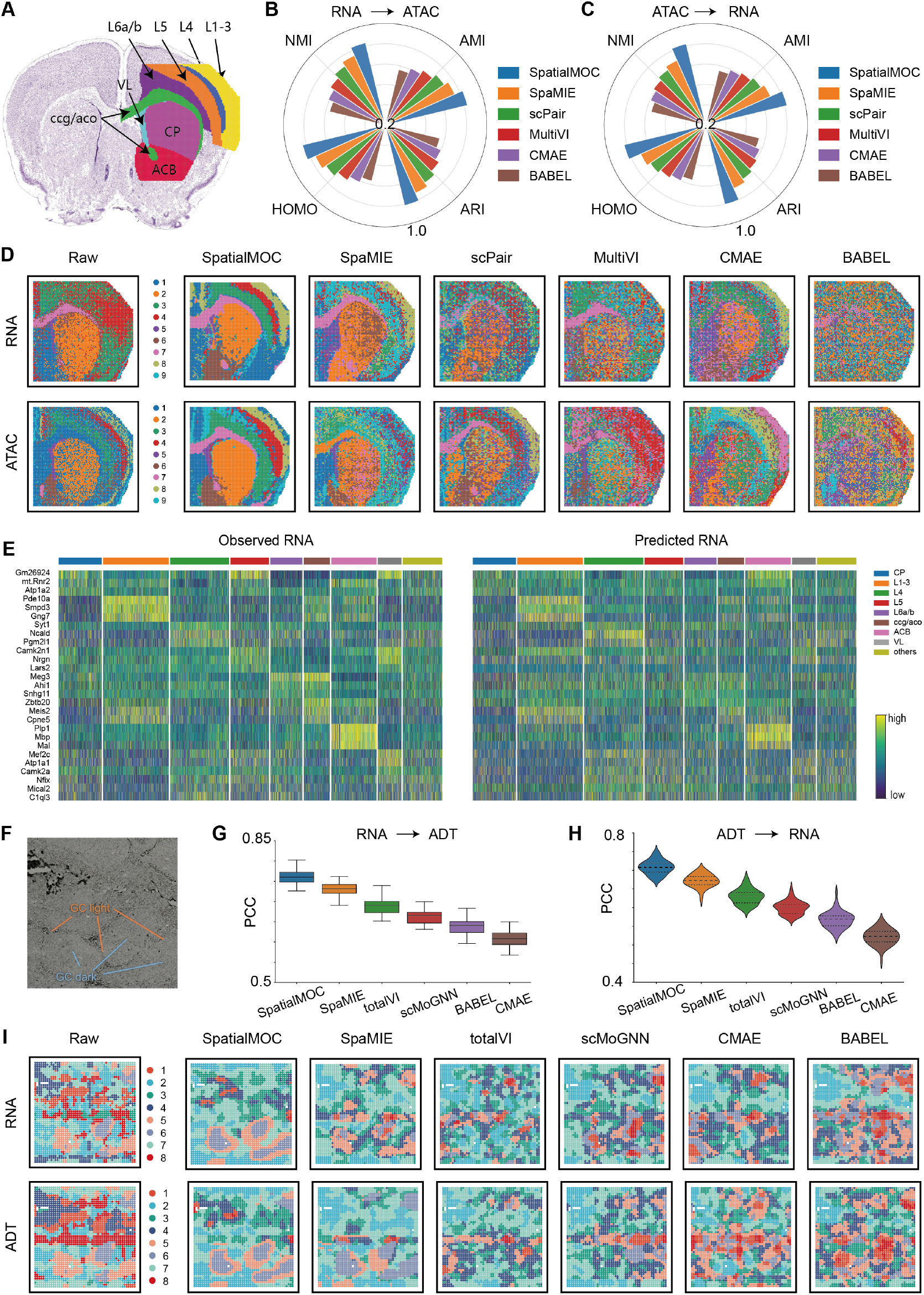
SpatialMOC faithfully preserves tissue architecture during cross-modality prediction. (**A**) Anatomical annotations of the mouse brain section. (**B–C**) Quantitative comparison of spatial domain identification following bidirectional RNA–ATAC prediction, evaluated using ARI, NMI, AMI, and HOMO. (**D**) Spatial domain reconstruction based on observed and predicted RNA and ATAC profiles across different computational methods. (**E**) Comparison of observed and reconstructed transcriptomic patterns across annotated anatomical regions. (**F**) H&E image of the human tonsil section showing the GC light and dark zones. (**G–H**) Quantitative comparison of bidirectional RNA–ADT prediction accuracy measured by PCC. (**I**) Spatial domain reconstruction from observed and predicted RNA and ADT profiles in the human tonsil.

Visual inspection further demonstrated that SpatialMOC faithfully reconstructed the spatial organization of the mouse brain (Figure 3D). Compared with existing methods, the predicted RNA and ATAC profiles exhibited improved spatial continuity and more clearly resolved anatomical boundaries. In particular, SpatialMOC accurately delineated multiple cortical layers (L1–3, L4, L5, and L6a/b) together with subcortical regions including the CP, ACB, ccg/aco, and ventrolateral nucleus (VL). Consistent with these spatial domains, reconstructed transcriptomic heatmaps closely matched the observed molecular patterns across annotated anatomical regions (Figure 3E). Together, these results indicate that SpatialMOC preserves both large-scale anatomical compartments and fine-scale cortical organization during cross-modality prediction.

We next assessed the generalizability of SpatialMOC using the Spatial-CITE-seq human tonsil dataset. The tissue exhibits well-defined germinal-center (GC) light and dark zones, providing a challenging benchmark for evaluating the preservation of fine-scale immune microenvironments (Figure 3F). SpatialMOC consistently achieved the highest prediction accuracy for both prediction directions. For RNA-to-ADT prediction, SpatialMOC reached a mean PCC of 0.7653, followed by SpaMIE, totalVI, and other baseline methods (Supplementary Table S11). Similar performance was observed for ADT-to-RNA prediction, where SpatialMOC achieved the highest mean PCC of 0.7124, followed by SpaMIE (Supplementary Table S12). The overall comparisons are shown in Figure 3G,H. Detailed quantitative results are summarized in Supplementary Tables S11 and S12.

The reconstructed molecular profiles also faithfully preserved the spatial organization of the human tonsil (Figure 3I). While several baseline methods produced fragmented spatial domains and blurred tissue boundaries, the RNA- and ADT-derived domains reconstructed by SpatialMOC accurately resolved the GC light and dark zones and closely matched both the observed molecular data and the underlying histological morphology. Collectively, these analyses demonstrate that SpatialMOC not only accurately predicts molecular profiles but also preserves the spatial organization required to reconstruct complex tissue architecture across distinct biological systems.

### SpatialMOC generalizes molecular profile prediction across biological and technical heterogeneity

To evaluate the generalizability of SpatialMOC beyond matched tissues and experimental settings, we assessed its performance across both biological and technical heterogeneity. Specifically, we performed cross-stage prediction using the MISAR-seq mouse embryo dataset (E15.5 and E18.5; spatial RNA and ATAC) and cross-platform prediction using two independently generated mouse spleen datasets, SPOTS and Spatial-CITE-seq (spatial RNA and ADT). These experiments provide stringent benchmarks for evaluating whether SpatialMOC can accurately reconstruct molecular profiles despite developmental progression, platform-specific variation, and heterogeneous signal distributions.

We first investigated whether SpatialMOC could generalize across developmental stages by training the model on embryonic day 15.5 (E15.5) tissues and predicting molecular profiles in embryonic day 18.5 (E18.5) tissues. The anatomical annotations of the E18.5 mouse embryo are shown in Figure 4A. Despite substantial developmental changes between the two stages, SpatialMOC consistently achieved the highest prediction accuracy. For ATAC-to-RNA prediction, SpatialMOC obtained the highest PCC (0.7221), followed by SpaMIE (0.6723) and scPair (0.6201) (Supplementary Table S14). The overall comparison is shown in Figure 4C. Similarly, SpatialMOC achieved an AUROC of 0.7513 for RNA-to-ATAC prediction (Figure 4B) together with the best 1 − FOSCTTM scores for both prediction directions (Figure 4D). Detailed forward(E15.5→E18.5) prediction results are summarized in Supplementary Tables S13 and S14. Reverse(E18.5→E15.5) prediction results are summarized in Supplementary Tables S19 and S20, with corresponding visual comparisons shown in Supplementary Figures S10 and S11.

**Fig. 4.**
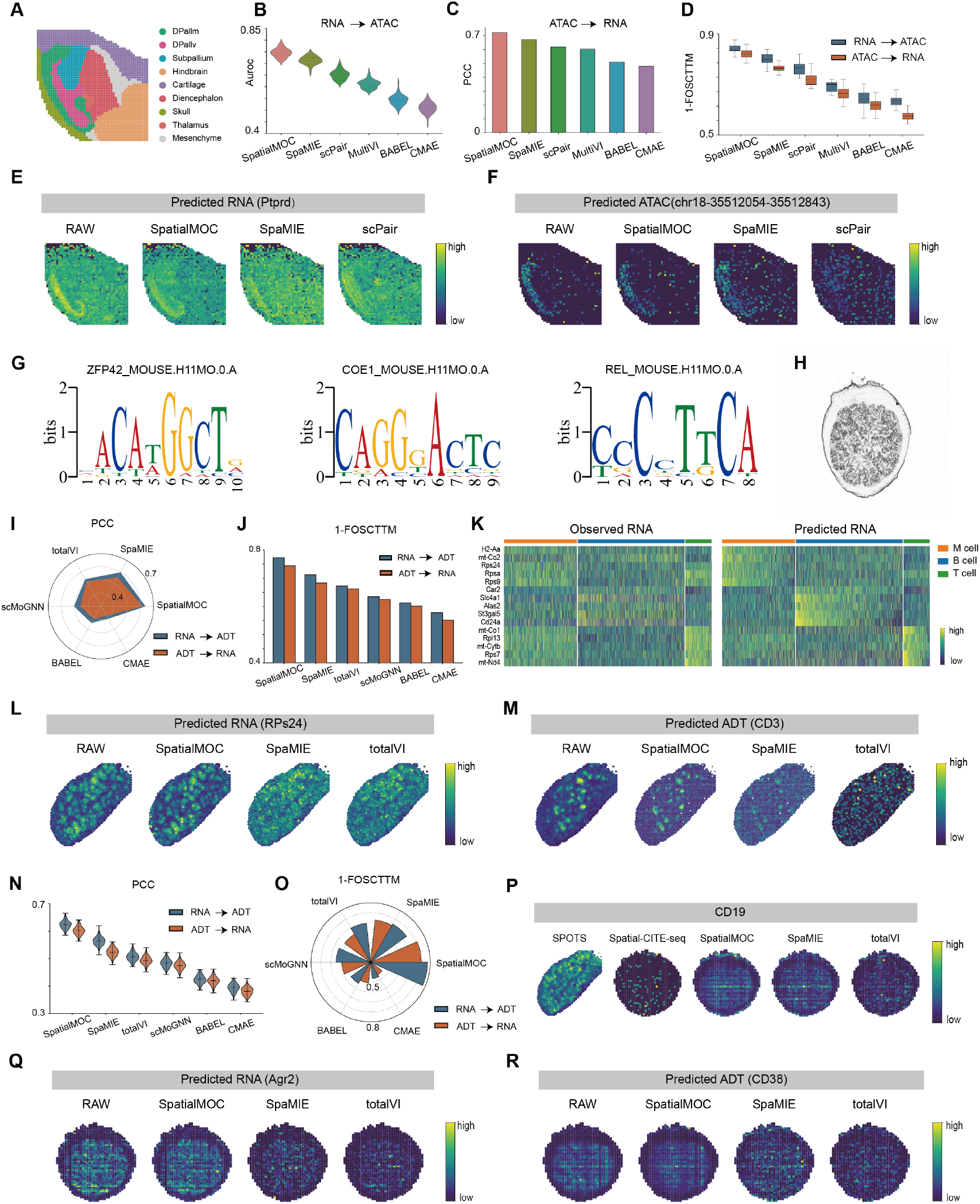
SpatialMOC generalizes across biological and technical heterogeneity. (**A**) Anatomical annotations of the E18.5 mouse embryo. (**B–D**) Quantitative evaluation of cross-stage prediction between E15.5 and E18.5: RNA-to-ATAC AUROC (**B**), ATAC-to-RNA PCC (**C**), and 1 − FOSCTTM for both directions (**D**). (**E–F**) Spatial reconstruction of the representative RNA marker *Ptprd* and ATAC peak chr18-35512054-35512843. (**G**) Motif enrichment analysis of reconstructed E18.5 ATAC profiles. (**H**) H&E image of the mouse spleen from the Spatial-CITE-seq dataset. (**I–J**) Quantitative evaluation of cross-platform prediction from Spatial-CITE-seq to SPOTS. (**K**) Comparison of observed and reconstructed transcriptomic patterns across immune cell clusters. (**L–M**) Spatial reconstruction of representative RNA (*Rps24*) and ADT (CD3) markers. (**N–O**) Quantitative evaluation of cross-platform prediction from SPOTS to Spatial-CITE-seq. (**P**) Recovery of the sparsely captured CD19 signal through cross-platform prediction. (**Q–R**) Spatial reconstruction of representative RNA (*Agr2*) and ADT (CD38) markers.

Beyond quantitative performance, SpatialMOC faithfully reconstructed developmentally regulated spatial molecular patterns. The predicted RNA expression of *Ptprd* accurately recapitulated its localized enrichment within the DPallm region (Figure 4E), while the predicted chromatin accessibility peak (chr18-35512054-35512843) reproduced the corresponding spatial accessibility landscape (Figure 4F). Motif enrichment analysis of the reconstructed E18.5 ATAC profiles further identified stage-specific transcription factor binding motifs, including ZFP42, COE1, and REL (Figure 4G), demonstrating that SpatialMOC preserves dynamic regulatory programs associated with developmental progression.

We next evaluated whether SpatialMOC could generalize across sequencing platforms using two independently generated mouse spleen datasets. Despite differences in experimental protocols and molecular capture efficiency, SpatialMOC consistently maintained superior prediction performance. The corresponding hematoxylin and eosin (H&E) staining image for Spatial-CITE-seq is shown in Figure 4H. When trained on Spatial-CITE-seq data to predict SPOTS molecular profiles, SpatialMOC achieved PCC values of 0.6334 for RNA-to-ADT prediction and 0.6123 for ADT-to-RNA prediction, followed by SpaMIE with the second-ranked values (Supplementary Tables S15 and S16). The overall comparison is shown in Figure 4I, together with the highest 1 − FOSCTTM scores (Figure 4J). Complete quantitative results are summarized in Supplementary Tables S15 and S16.

The reconstructed transcriptomic profiles accurately preserved the spatial organization of immune cell populations (Figure 4K). Moreover, SpatialMOC faithfully recovered complex spatial expression patterns across heterogeneous immune compartments. The predicted RNA expression of *Rps24* reproduced its broad yet regionally enriched distribution, whereas the predicted ADT signal of the T-cell marker CD3 accurately delineated the periarteriolar lymphoid sheath (PALS) within the white pulp (Figure 4L,M). Compared with baseline methods, SpatialMOC generated substantially sharper spatial boundaries, indicating improved reconstruction of highly localized immune microenvironments.

Finally, we evaluated the robustness of SpatialMOC under platform-specific signal dropout by reversing the prediction direction and reconstructing Spatial-CITE-seq profiles using SPOTS as the reference. SpatialMOC again achieved the highest PCC and 1 − FOSCTTM scores (Figure 4N,O), with complete quantitative results provided in Supplementary Tables S17 and S18. Importantly, SpatialMOC successfully recovered biologically meaningful molecular signals that were sparsely captured in the original Spatial-CITE-seq data. For example, the reconstructed CD19 distribution accurately identified B-cell follicles within the white pulp despite the low detection efficiency of the raw measurements (Figure 4P). Likewise, the predicted expression patterns of *Agr2* and CD38 faithfully reproduced their compartment-specific spatial organization (Figure 4Q,R). Collectively, these results demonstrate that SpatialMOC remains robust across developmental progression, sequencing platforms, and heterogeneous signal quality.

### SpatialMOC robustly reconstructs spatial molecular landscapes from degraded measurements

To evaluate the robustness of SpatialMOC under technically compromised conditions, we investigated its ability to reconstruct complete spatial molecular profiles from degraded measurements containing simulated signal dropout. Cross-slice masking experiments were performed using the MISAR-seq mouse embryo E15.5 dataset (spatial RNA and ATAC) and the Stereo-CITE-seq mouse thymus dataset (spatial RNA and ADT). For each dataset, increasing levels of masking (0–50%) were introduced into the input modality, and SpatialMOC was used to reconstruct the complete target molecular profiles. These experiments provide a stringent assessment of the model’s robustness to technical noise and incomplete spatial measurements.

We first evaluated spatial interpolation using the mouse embryo dataset. Across all masking ratios, SpatialMOC consistently maintained the highest prediction accuracy. In the RNA-to-ATAC task, SpatialMOC achieved a PCC of 0.7646 even when 20% of the input signals were masked, followed by SpaMIE (0.7394) and scPair (0.6907) (Supplementary Table S21). The overall masking trajectory is shown in Figure 5A. Similar robustness was observed for ATAC-to-RNA prediction (Figure 5B). Complete quantitative results across masking rates from 0% to 50% are summarized in Supplementary Tables S21 and S22. Because a 20% masking rate represents a realistic level of technical signal loss, we used this condition for detailed downstream evaluation. Under this setting, SpatialMOC achieved the highest median 1 − FOSCTTM scores (0.8431 for RNA-to-ATAC and 0.8171 for ATAC-to-RNA), indicating that biologically meaningful molecular relationships were largely preserved despite substantial information loss (Figure 5C). Detailed interpolation results under the 20% masking condition are provided in Supplementary Tables S29 and S30.

**Fig. 5.**
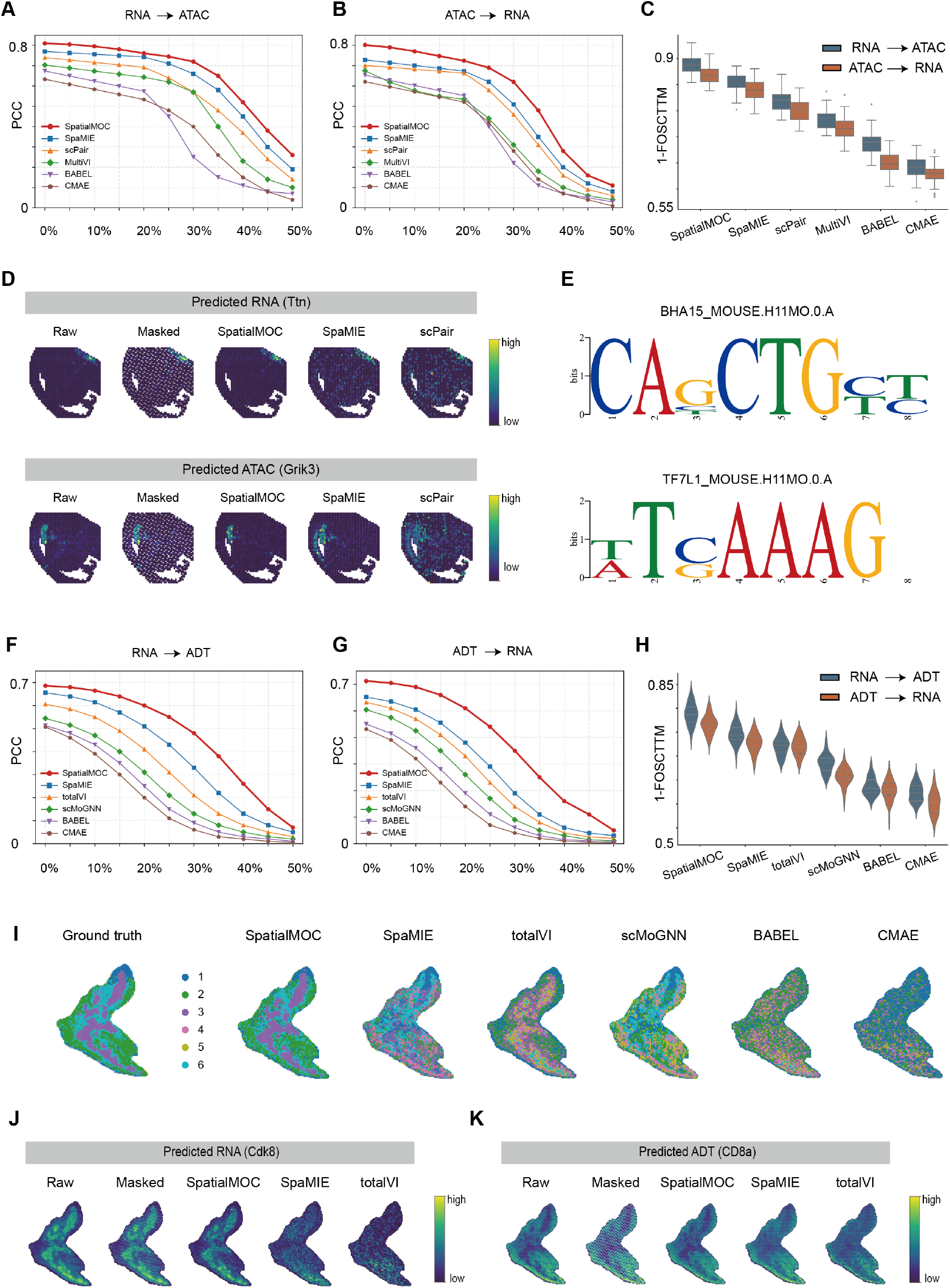
SpatialMOC robustly reconstructs spatial molecular landscapes from degraded measurements. (**A–B**) Prediction accuracy across increasing masking ratios (0–50%) for bidirectional RNA–ATAC interpolation in the mouse embryo dataset. (**C**) Comparison of 1 − FOSCTTM scores under the representative 20% masking condition. (**D**) Spatial reconstruction of the representative RNA marker *Ttn* and ATAC marker *Grik3* from degraded measurements. (**E**) Motif enrichment analysis of reconstructed ATAC profiles following spatial interpolation. (**F–G**) Prediction accuracy across increasing masking ratios for bidirectional RNA–ADT interpolation in the mouse thymus dataset. (**H**) Comparison of 1 − FOSCTTM scores under the representative 20% masking condition. (**I**) Recovery of thymic tissue architecture based on reconstructed ADT profiles under the 20% masking condition. (**J–K**) Spatial reconstruction of representative RNA (*Cdk8*) and ADT (CD8a) markers in the mouse thymus.

Beyond quantitative performance, SpatialMOC faithfully recovered spatially localized molecular patterns from degraded inputs. At the 20% masking level, the reconstructed RNA expression of *Ttn* accurately reproduced its enrichment within developing muscle, while the predicted chromatin accessibility of *Grik3* correctly recovered its spatial localization in the upper DPallm and subpallium (Figure 5D). Motif enrichment analysis of the reconstructed ATAC profiles further recovered biologically relevant transcription factor binding motifs, including BHA15 and TF7L1 (Figure 5E), demonstrating that SpatialMOC preserves regulatory information despite substantial technical signal loss.

We next evaluated SpatialMOC on spatial transcriptomic and proteomic data using the mouse thymus dataset. Across all simulated masking ratios, SpatialMOC consistently achieved the highest prediction accuracy for both RNA-to-ADT and ADT-to-RNA interpolation (Figure 5F,G). Complete quantitative evaluations are summarized in Supplementary Tables S23 and S24. Under the representative 20% masking condition, SpatialMOC achieved a PCC of 0.5951 for RNA-to-ADT prediction, followed by SpaMIE (0.5054) and totalVI (0.4068) (Supplementary Table S23), while maintaining the highest 1 − FOSCTTM scores for both prediction directions (Figure 5H; Supplementary Tables S31 and S32).

SpatialMOC also accurately reconstructed thymic tissue architecture from degraded measurements. Spatial clustering of the reconstructed ADT profiles accurately recovered the lobular organization of the thymus and closely matched the ground-truth tissue structure, whereas competing methods produced fragmented domains and blurred spatial boundaries (Figure 5I). Consistent with these structural improvements, SpatialMOC correctly reconstructed the restricted inner medullary expression of *Cdk8* together with the broad cortical distribution of the ADT marker CD8a (Figure 5J,K). These results demonstrate that SpatialMOC effectively restores both molecular signals and higher-order tissue architecture from degraded spatial measurements.

Finally, we evaluated the robustness of SpatialMOC by reversing the reference and target tissue sections for spatial interpolation. SpatialMOC again consistently achieved the best performance across all evaluation metrics, indicating that its interpolation accuracy is independent of prediction direction. The complete reverse-direction masking trajectories are provided in Supplementary Tables S25–S28, and the detailed results under the 20% masking condition are provided in Supplementary Tables S33–S36. Corresponding visual comparisons are shown in Supplementary Figures S12–S15.

## Discussion

Spatial multi-omics technologies are transforming our ability to investigate tissue organization by simultaneously characterizing complementary molecular layers within their native spatial context. However, the widespread adoption of these technologies remains constrained by the high cost, limited throughput, and technical challenges associated with generating paired multi-omics measurements. In this study, we developed SpatialMOC, a computational framework that reconstructs missing spatial molecular modalities from available measurements while explicitly preserving spatial organization. Across multiple tissues, molecular modalities, developmental stages, sequencing platforms, and levels of data quality, SpatialMOC consistently demonstrated accurate cross-modality prediction together with faithful reconstruction of biologically meaningful tissue architecture. These results highlight the potential of computational prediction to complement experimental spatial multi-omics profiling and expand the utility of existing single-omics datasets.

A key strength of SpatialMOC lies in its explicit integration of spatial context with cross-modality representation learning. Unlike existing computational approaches that primarily focus on molecular correspondence, SpatialMOC jointly models tissue topology and molecular relationships, enabling the reconstruction of spatial molecular landscapes rather than isolated molecular features. Consequently, the predicted profiles preserve not only quantitative molecular signals but also higher-order biological organization, including cortical lamination in the developing brain, immune compartmentalization in the spleen and tonsil, and tissue architecture under substantial technical signal loss. The ability to maintain these biologically meaningful spatial patterns is particularly important for downstream analyses such as spatial domain identification, tissue annotation, and the investigation of cellular microenvironments.

Our results further demonstrate that SpatialMOC generalizes across diverse sources of biological and technical heterogeneity. The framework accurately predicted molecular profiles across developmental stages, sequencing platforms, and degraded datasets with substantial simulated dropout, indicating that the learned representations capture fundamental spatial-molecular relationships that are robust to experimental variation. More broadly, methods for integrating heterogeneous datasets and mapping new observations to reference atlases have demonstrated the value of representations that remain stable across batches, platforms, and study-specific variation [38, 39]. As spatial transcriptomics technologies continue to diversify, computational frameworks capable of integrating heterogeneous datasets and recovering missing molecular information will become increasingly valuable for constructing comprehensive spatial atlases and enabling large-scale comparative analyses across tissues, developmental processes, and disease states.

Several limitations should also be acknowledged. First, the current implementation focuses on two-dimensional spatial datasets, whereas extending the framework to three-dimensional spatial reconstruction could provide a more comprehensive representation of tissue organization. Second, SpatialMOC currently relies on paired multi-omics reference data for model training. Developing strategies that reduce this dependency, such as semi-supervised or self-supervised learning, may further broaden its applicability. Third, as spatial technologies continue to advance toward single-cell and subcellular resolution, additional methodological improvements will be required to accommodate increasingly sparse measurements and larger datasets. Finally, extending the framework to incorporate emerging molecular modalities, including spatial metabolomics and lipidomics, may enable more comprehensive characterization of tissue regulatory networks.

In summary, SpatialMOC provides a unified computational framework for reconstructing spatial multi-omics landscapes from incomplete molecular measurements. By combining accurate cross-modality prediction with preservation of tissue architecture and robustness across heterogeneous biological settings, SpatialMOC extends the analytical potential of existing spatial omics datasets and provides a practical foundation for future studies of tissue organization, development, and disease.

## Acknowledgments

J.C., T.W. and Z.X.L. conceived the study and established the research plan. J.C. and Z.X.L. were responsible for designing the core research framework. P.M.Z. and B.T.W. designed the auxiliary experimental framework and relevant optimization strategies. H.S. and Y.Z. were in charge of collecting and sorting the research-related datasets, ensuring the validity and completeness of the data. H.S., Y.Z., K.Y.Z., Y.T.W. and J.L.H. collected the benchmarking method scripts, carried out experimental verification, and generated the primary experimental results. J.C., T.W. and Z.X.L. analyzed the experimental results, summarized the research conclusions, and prepared and organized the figures and tables in the manuscript. J.C., T.W. and Z.X.L. wrote the initial draft of the manuscript, and all authors participated in the revision and polishing of the manuscript content. J.C., T.W. and Z.X.L. supervised the entire research process, including experimental design, result analysis and manuscript revision. T.W. only participated in this study during his affiliation with Northwestern Polytechnical University. All authors read and approved the final manuscript.

This work was supported by the National Natural Science Foundation of China (grant numbers: 62402382, 62572391) and the National Key R&D Program of China (grant number: 2025YFC3410200).

